# Gallic Acid, a Methyl 3,4-Dihydroxybenzoate Derivative, Induces Neural Stem Cells to Differentiate and Proliferate

**DOI:** 10.1101/2021.03.01.433474

**Authors:** Junxing Jiang, Weiyi Liu, Jitao Hai, Yan Luo, Keqi Chen, Yirong Xin, Junping Pan, Yang Hu, Qin Gao, Fei Xiao, Huanmin Luo

## Abstract

Differentiation and proliferation of neural stem cells (NSCs) are both important biological processes in cerebral neural network. However, these two capacities of NSCs are limited. Thus, the induction of differentiation and/or proliferation via administration of small molecules derived from natural plants can be considered as a potential approach to repair damaged neural networks. This paper reports that gallic acid, a catechol compound selected from derivatives of methyl 3,4-dihydroxybenzoate (MDHB), selectively induces NSCs to differentiate into immature neurons and promotes proliferation by activating phosphorylation of key proteins in the MAPK-ERK pathway. In addition, we found that 3,4-dihydroxybenzoic acid was the main active structure which could show neurotrophic activity. The substitution of carboxyl group on the benzene ring into ester group may promote differentiation on the basis of the structure of 3,4-dihydroxybenzoic acid. Meanwhile, the introduction of 5-hydroxyl group may promote proliferation. Generally, this study identified a natural catechol compound that promotes differentiation and proliferation of NSCs in vitro.

## 1. Introduction

Neural stem cells (NSCs) are cells in the nervous system which can self-renew and differentiate into a variety of specific nerve cell types. Usually, NSCs divide in two ways: symmetrical cell division and asymmetric cell division[1,2]. Symmetrical cell division refers to the division of a NSCs into two daughter cells identical to a metrocyte, while asymmetric cell division is a common form of NSCs division after brain tissue is stimulated by external factors such as injury. The balance of these two forms can keep the stem cell bank in a stable state under normal physiological conditions[3]. Therefore, to induce the differentiation or proliferation of NSCs under certain conditions by virtue of this characteristic can be regarded as a new idea for the treatment in neurodegenerative diseases such as Alzheimer’s disease (AD) and Parkinson’s disease (PD).

Small molecule components extracted from natural plants show strong biological activity in neurocyte. Methyl 3,4-dihydroxybenzoate (MDHB, C_8_H_8_O_4_) is a natural small molecule compound with a molecular weight of 168.147 (CAS), which mainly exists in traditional herbs such as *Thalictrum thunbergi DC* and *Aster indicus L*. Our laboratory has carried out researches on MDHB for many years. In the previous studies, we found that MDHB had 3,4-dihydroxyl (catechol, C_6_H_6_O_2_) structure and a variety of activities such as antioxidation, anti-neurotoxicity, neurotrophy and neuroprotection[4-6]. Especially, MDHB can promote the growth of primary neuronal processes and the differentiation of NSCs into cholinergic neurons in vitro by inhibiting Akt phosphorylation and activating autophosphorylation of GSK3β at tyrosine 216[7]. Besides, some other compounds with polyphenol structure have also been shown to have neurotrophic and antioxidant and antibacterial effects. For example, curcumin can quickly enter the brain through the blood-brain barrier and preferentially bind to Aβ plaques, effectively inhibiting the enlargement and aggregation of Aβ plaques[8,9]. Resveratrol may inhibit the formation of inducible nitric oxide synthase (iNOS) and pro-inflammatory mediators of hippocampal neurons in vitro. In the hypoxia injury mode, the antioxidant capacity of microglia cells can also be improved[10]. Quercetin was been proved that it can reduce ROS production, increase MnSOD activity, reduce mitochondrial swelling and chromatin coagulation induced by aluminum in the alum-induced mitochondrial oxidative injury SD rat model, and finally protect the degenerated neurons from death[11]. However, whether these polyphenols have biological activities that promote the differentiation or proliferation of NSCs has not been investigated.

We have recently preliminarily screened a series of polyphenols from MDHB derivatives. Some of these compounds have been proved to have antibacterial, antioxidant and neurotrophic activities[12-15], but whether they have differentiated or even proliferative activities in NSCs remains to be explored. In the current study, we found that gallic acid, a derivative of MDHB, can induce the differentiation and proliferation of NSCs. Besides, it also plays a proliferative role by activating the phosphorylation of key protein ERK1/2 in the downstream MAPK-ERK pathway of PDGF receptor.

## 2. Materials and methods

### 2.1 Animals

This study was carried out in accordance with the recommendations of the Animal Research committee of the School of Medicine of Jinan University. The experimental procedures were approved by the Institutional Animal Care and Use Committee of Institute of Laboratory Animal Science, Jinan University.

### 2.2 *In Vitro* Culture of Rat Primary Neural Stem Cells

The methods of NSCs isolation and culture were based on the other published article[7,16]. In brief, the newborn rat hippocampi (< 24 h) were chopped and digested by 0.25% trypsin (Gibco) for 8 min in 5% CO_2_ incubator at 37°C. Equal volume of DMEM/F12 medium (Gibco) containing 10% fetal bovine serum (Lonsera) was added to the cell suspension, then the cell suspension was filtered by 70 µm microfiltration membrane and centrifuged for 5 min. The cells which were then resuspended in DMEM/F12 supplemented with 20 ng/mL EGF (Proteintech), 10 ng/mL basic fibroblast growth factor (Proteintech), 2% B27 without vitamin A, and 1% Penicillin-Streptomycin (Sigma), were seeded in 6 well plate at a density of 10,000 cells per well and cultured in 5% CO_2_ incubator at 37°C for about 5 days. Within these days, the cells grew into suspended neurospheres which could be gathered and passaged via mechanical pipette method.

### 2.3 NSCs Differentiation and Proliferation

Neurospheres which suspension cultured to the fifth day were collected and centrifugated for 5 min, then dissociated into single cells by stem cell acctuse (STEM CELL) and seeded in glass cover slips coated by 0.0125 mg/mL poly-D-lysine (PDL, Sigma) at the density of 50,000 cells/mL in DMEM/F12 medium containing 1% FBS (Lonsera) and 1% penicillin and streptomycin. When cells completely adherent after 2h, the original culture medium was replaced by neural-basal medium (NeuroCult), which containing different concentration of protocatechuic acid (PCA), vanillic acid (VA), gallic acid (GA), methyl 2,3-dihydroxybenzoate, methyl 2,5-dihydroxybenzoate, methyl 3,5-dihydroxybenzoate, methyl vanillate (MV), methyl gallate (MG), and MDHB at 0, 1, 5, 10, 20, 25, 30, and 40 µM, respectively.

### 2.4 Cell Counting Kit – 8 Assay

Cells viability was evaluated by were cultured in 96-well culture plate in different concentrations of drug medium respectively 1h, 2h, 3h, and 4h. 10µL CCK8 solution (Dojindo) was added to each well and the absorbance was analyzed at 450 nm by Multiskan Sky Microplate (PerkinElmer). Cell viability rate was calculated based on the product specification.

### 2.5 Immunofluorescence Assay

Cells were fixed in 4% paraformaldehyde (PFA), washed with phosphate-buffered saline (PBS, pH7.6) and blocked with pre-cooled super blocking buffer for 1h at RT which containing 1% fish serum, 0.5% goat serum 0.5% donkey serum and 0.5% bull serum albumin dissolved in 0.3% PBST. Afterwards, the cells were incubated with the primary antibodies of interested proteins at 4°C overnight (Nestin, 1:1000, Sigma; Tuj1, 1:1000, Sigma; GFAP, 1:1000, Abcam; Ki67, 1:1000, ProteinTech). The next day, the cells were washed with 0.3% PBST for three times and incubated with immunofluorescence secondary antibodies for 2h at RT in darkness (Cy3, 1:1000, Earthox; 488, 1:1000, Earthox). 4’,6-diamidino-2-phenylindole (DAPI, 1:1000, Sigma) was used to show nuclei clear and dyed for 5 min. The signals from the Immunofluorescence staining cells were then captured by a confocal microscope (Zeiss, LSM700). Finally, the photographs were analyzed via Image J software while the Nestin, Tuj1, GFAP positive cell numbers and cell nuclei counted in each of ten random fields per well.

### 2.6 Western blot

The cell protein expression levels of PH3, Akt, p-Akt, ERK1/2, and p-ERK, after compounds treatment for 5d, were detected by Western blot. The preparation of cell lysates, western blotting and detection of corresponding immunoreactive protein bands were performed as described in previous reports in our laboratory[7,17]. The samples were incubated with the following primary antibodies: β-actin (1:1000, CST), PH3 (1:1000, CST), Akt (1:1000, Sigma), p-Akt (1:1000, Ser10, CST), ERK1/2 (1:1000, CST), and p-ERK (1:1000, Thr202/Tyr204, CST). Finally, the membrane was washed and submerged in chemiluminescent HRP substrate while the fluorescent signals of the blots were captured by gel imaging analysis system (Bio-rad). The membrane was incubated with primary antibody of β-actin (1:1000, CST). The following steps repeated the above except the secondary antibody replaced to HRP-conjugated goat IgG against rabbit (1:2000, Earthox).

### 2.7 mRNA Collection and Transcriptome Analysis of GA-Induced Proliferation

Cells were respectively collected after treated GA for 24h, 48h, 72h and 5d. Total mRNA were extracted using the pre-cooled RNase-free TRIzol reagent (Invitrogen), subsequently, the concentration and purity were determined using the Nanodrop 2000C microvolume spectrophotometer (Thermo Fisher). Following the manufacturer’s protocol, the cDNA fragments were purified using a PCR extraction kit and enriched by PCR to constructed cDNA library. RNA-seq technology was used to analyze gene expression in GA-induced NSCs proliferation.

### 2.8 Statistical Analysis

Statistical analysis of data, which were presented as mean ± S.E.M, were performed by using a *t*-test to compare two independent sample groups, and one-way ANOVA to compare three or more sample groups. *P* < 0.05 was considered as statistically significant.

## 3. Results

### 3.1 Culture and Identification of Rat Primary NSCs

Primary NSCs derived from suspension cultured neurospheres in vitro. When passed to the second generation, the number and size of the neurospheres were significantly increased (**Fig. 1B**). Nestin and DAPI were used for immunofluorescence staining of neurospheres (**Fig. 1A**). The result showed that more than 80% cells were identified as Nestin positive cells (**Fig. 1C**).

**Fig. 1.**
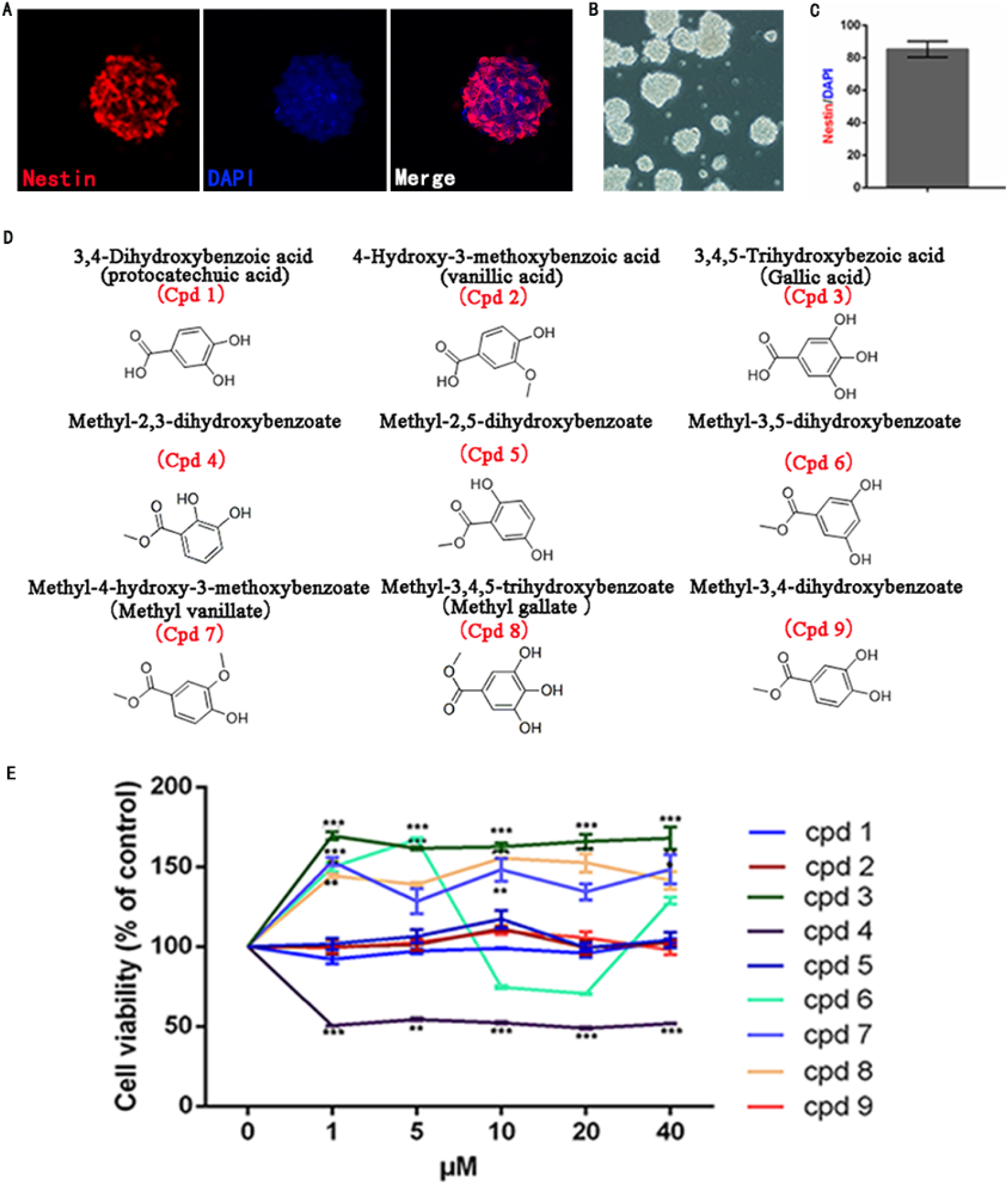
The morphological and purity identification of primary neurospheres and NSCs and effects of MDHB derivatives on the viability of NSCs. **(A)** Nestin and DAPI fluorescent staining of primary neurospheres consisted of NSCs. **(B)** The morphology of neurospheres in vitro. **(C)** The percentage of Nestin and DAPI of neurospheres. **(D)** The structural formula of MDHB and 8 candidate compounds. **(E)** Cell viability results of NSCs treated by Cpd1, 3, 7, 8, and 9 at 5, 10, 20, and 40 µM dose for 4h, respectively. A dose of 0 µM indicates the treatment of DMSO as a solvent control. (^*^*P*_<_0.05, ^**^*P*_<_0.01, ^***^*P*_<_0. 001, compared with DMSO group, ^#^*P*_<_0.05, ^##^*P* _<_0.01, ^###^*P*_<_0. 001, compared with MDHB group, n=3).

### 3.2 Cell Viability of MDHB Derivatives on NSCs

As shown in **(Fig. 1D)**, we have selected 8 compounds which were based on the structural similarity with MDHB and different from it by only one group: PCA (Cpd1), VA (Cpd 2), GA (Cpd3), methyl 2,3-dihydroxybenzoate (Cpd4), methyl 2,5-dihydroxybenzoate (Cpd5), methyl 3,5-dihydroxybenzoate (Cpd6), MV (Cpd7), and MG (Cpd8), and MDHB was Cp d9 **(Fig. 1D)**. Most of these candidate compounds come from natural plants and have the basic structure of catechol. The neurospheres were digested and inoculated in 96-well plate s. After 2h, the cells were adhering and cultured in drug-containing medium and the cell viability was measured using CCK-8 kit. The concentration gradient of drug containing medium was set as: 0, 1, 5, 10, 20, and 40µM. All of the compounds except Cpd4 had lower toxicity on NSCs **(Fig. 1E)**. Therefore, we screened the top5 compounds least toxicity on NSCs, which were PCA (Cpd1), GA (Cpd3), MV (Cpd7), MG (Cpd8), and MDHB (C pd9), from these derivatives for the next experiments.

### 3.3 MG, GA, and MDHB promote the differentiation of NSCs into immature neurons

Considering the effect of MDHB on the differentiation of NSCs into neuronal cells described in previous studies, we determined whether the preliminarily screened compounds could induce NSCs differentiation to neurons. The neurospheres were digested into single NSCs and treated Cpd1, 3, 7, 8, and 9 at the concentration of 0, 5, 10, 20, and 40 µM, respectively. After five days, it could be seen that cells in each group showed different morphological characteristics. After statistical analysis of cells with neuronal morphology using Image J software, we found that Cpd8 (MG), Cpd3 (GA), and Cpd9 (MDHB) induced more immature neurons within the concentration range from 20 to 40 µM **(Fig. 2B)**. To further determine the optimum concentration of these compounds, we treated these candidates at 25 µM and 30 µM. **(Fig. 2C)**. As shown in **(Fig.2A**) the differentiation effects of these 3 compounds on NSCs was the most significant at 25 µM, also, the number of cells in GA group showed the most significant neuronal morphology.

**Fig. 2.**
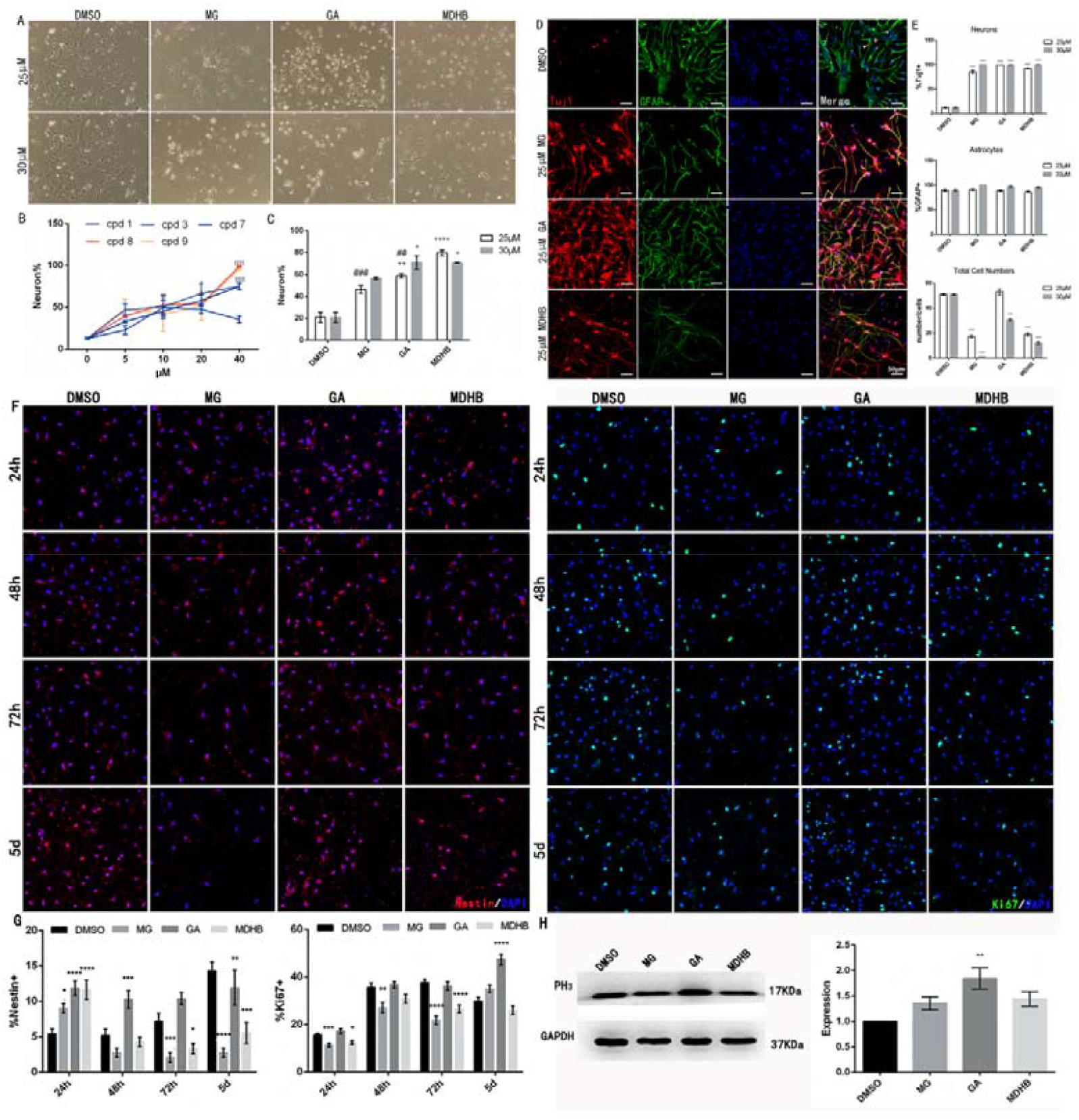
Differentiation and proliferation effects of MG, GA, and MDHB on NSCs. **(A)** Morphological analysis of the cells differentiated from NSCs treated by MG, GA and MDHB. **(B)** Statistical analysis of differentiation rate of NSCs treated by Cpd1, 3, 7, 8, and 9 at 5, 10, 20, and 40 µM, respectively. **(C)** Statistical analysis of differentiation rate of NSCs treated by MG, GA and MDHB at 25 and 30 µM, respectively (^*\**^*P*_<_0.05, ^*\*\**^*P*_<_0.01, ^*\*\*\*\**^*P*_<_0.0001, compared with DMSO group, ^*##*^*P* _<_ 0.01, ^*####*^*P* _<_ 0.0001, compared with MDHB group, n=3). **(D)** Immunostaining of cells differentiated by MG, GA, and MDHB-induced NSCs using the neuron marker Tuj1 (red dyed) and the astrocyte marker GFAP (green dyed). DAPI is the staining of nucleus (blue dyed). **(E)** Statistical analysis of immature neurons differentiated, astrocytes differentiated, and total cell numbers by MG, GA, and MDHB-induced NSCs. (^****^*P*_<_0.0001, compared with DMSO group, n=3). (F) Immunostaining of cells proliferated by MG, GA, and MDHB-induced NSCs by using the cell proliferation-related marker Ki67 and the NSCs marker Nestin. (G) Statistical analysis of cells proliferated by MG, GA, and MDHB-induced NSCs. (H) Western blot was used to analyze the proteins from cells treated by MDHB derivatives. (^****^*P*_<_ 0.0001, compared with DMSO group, n=3).

Neuronal marker Tuj1 and astrocyte marker GFAP are commonly used to identify cell types. Tuj1 (immature neurons) and GFAP (astrocytes) were detected by cellular immunofluorescence technology **(Fig. 2D)**, and the results showed that MG, GA, and MDHB could promote the expression of Tuj1 in NSCs, and GA group was the most significant **(Fig. 2E)**. Likewise, the number of cells in GA group was significantly higher than that in other two treatment groups **(Fig. 2E)**.

### 3.4 Induction of proliferation of GA on NSCs

Given our results demonstrating that the number of cells in GA group was significantly increased, we next questioned whether GA itself, MG, or MDHB played an important role in promoting proliferation. To verify this, NSCs marker Nestin and mitotic marker Ki67 were used for immunofluorescence staining. Additionally, the expression levels of cell cycle marker PH3 were determined by Western blot in order to further elucidate the proliferative effects after prolonged administration time of the 3 candidates on NSCs. PH3 exists in late G2 period and M period which is widely used to mark proliferating and dividing cells in hippocampus. Immunofluorescence results showed that the protein expression levels of Nestin and Ki67 in GA group increased significantly after the 3 candidates treated for 24h, 48h, 72h, and 5d, respectively **(Fig. 2F,G)**. Analysis of Western blot confirmed that protein expression of PH3 was increased in GA group. Instead, it was basically unchanged in both MG and MDHB groups **(Fig. 2H)**. We therefore focused on GA in subsequent studies.

### 3.5 Transcriptome Sequencing Analysis of the Proliferation Promotion Effects of GA in NSCs

According to above phenomenon that GA can induce NSCs to differentiate into immature neurons and continuously induce proliferation, we then explored the potential proliferation mechanism. The mRNA of NSCs administrated by 25µM GA was collected for transcriptome sequencing analysis. We found that 849 genes were significantly upregulated as well as 79 genes were significantly down-regulated in the GA group **(Fig. 3A)**, Furthermore, GA may mainly act on the PDGF receptor located on the cell membrane of NSCs in the calcium signaling pathway to promote proliferation. **(Fig. 3B)**.

**Fig. 3.**
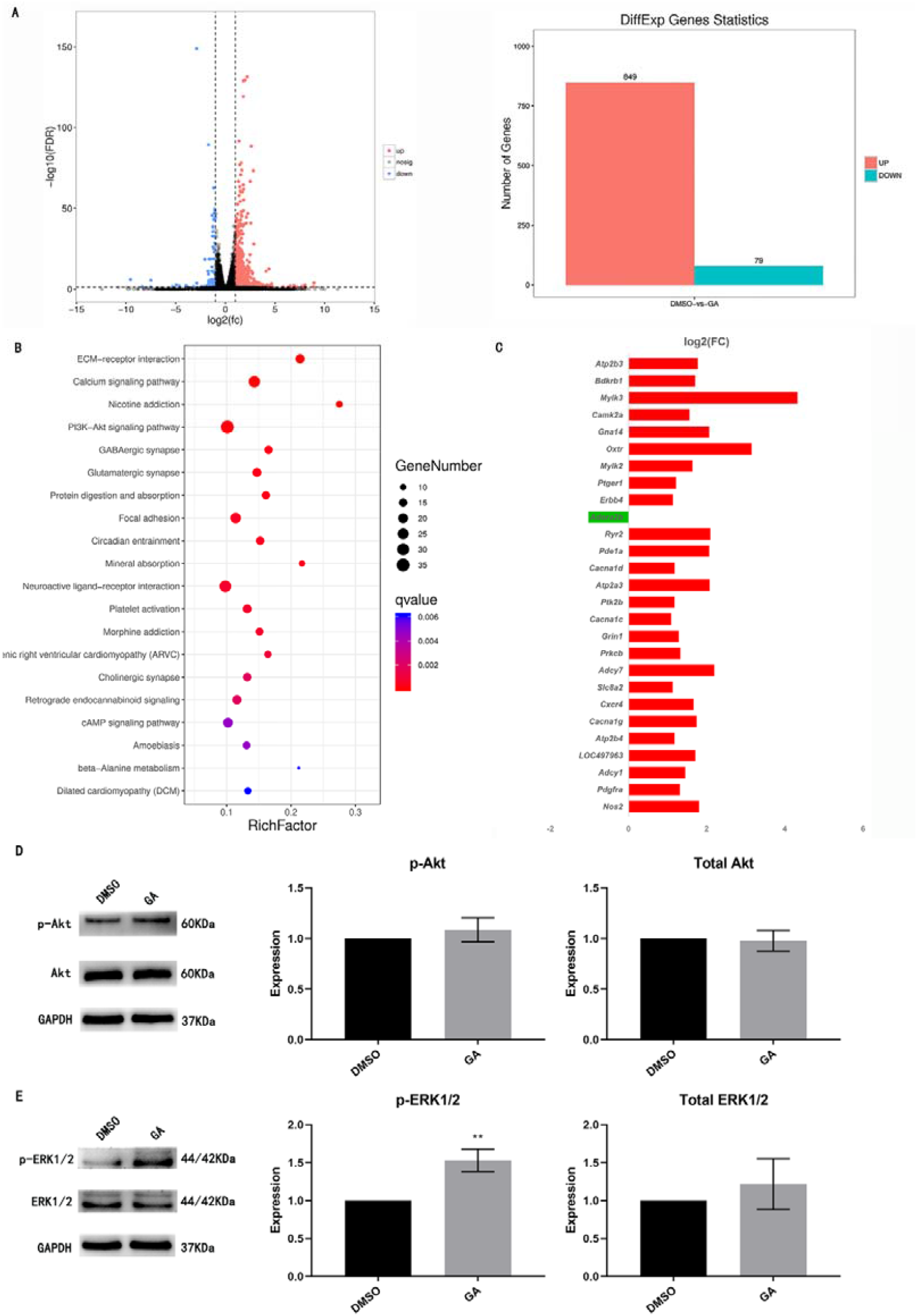
Transcriptomic analysis of NSCs proliferation induced by GA and effects of GA on Akt and ERK1/2 proteins on signal transduction pathways. **(A)** Statistical result of all differentially expressed genes. **(B)** Top 20 of KEGG pathway enrichment of NSCs proliferation induced by GA. **(C)** Sequencing analysis of all differential genes in calcium signaling pathway. **(D, E)** Western blot was used to analyze Akt and ERK1/2 proteins from cells treated by GA. The Akt and ERK1/2protein blots obtained by electrophoresis were compared and quantified with GAPDH, which is the intrinsic reference in the sample. (^*^*P* < 0.05, n=3).

### 3.6 Effects of GA on the phosphorylation level of Akt and ERK1/2 in NSCs

There are several signal transduction pathways downstream of PDGF receptor. Combining the theoretical analysis and above sequencing results, we hypothesized that GA may conduct downstream signal transduction through PI3K-Akt or MAPK-ERK pathways. Western blot was used to detected the phosphorylation levels of Akt and ERK1/2 these were the key proteins of these two pathways **(Fig. 3D,E)**. Our data revealed that the expressions of phosphorylated Akt protein and the total Akt protein in GA group remained unchanged compared with the control group after 25 µM GA administrated. However, the phosphorylation level of ERK1/2 at threonine 202 (Thr202) and tyrosine 204 (Tyr204) was upregulated while the expression level of total ERK1/2 not changed **(Fig. 3D,E)**. Thus, GA may induce proliferation by activating phosphorylation of ERK1/2 protein in the MAPK-ERK pathway.

## 4. Discussion

In this study, we screened 8 derivatives of MDHB which contain polyphenol structure and used MDHB as the control drug, and identified 3 compounds (MG, GA, and MDHB) with differentiation effect in NSCs. Also, GA has proliferation promotion effect in NSCs, which is achieved by activating MAPK-ERK pathway. Considering the differentiation and proliferation of NSCs are complex physiological processes, the exploration of the effects of pharmacological manipulations such as the use of traditional drugs on the growth and proliferation of NSCs may provide new ideas for the development of stem cell therapies of neurodegenerative diseases.

In recent years, many studies have shown that NSCs can simultaneously proliferate and differentiate into neurons under the same stimulation[18-22], but whether MDHB and its derivatives have both these two effects is unclear. In this study, we prepared rat hippocampal NSCs with high purity, and confirmed that MG, GA, and MDHB of 9 tested compounds have the ability to induce the differentiation of NSCs into neurons. Furthermore, GA can also induce the proliferation of NSCs. These results clearly demonstrate the effects of GA on NSCs in vitro. Therefore, we hypothesized that GA may change the ratio of differentiation and proliferation, making the proliferation promotion effect equally significant, while the proportion of differentiated cells was generally higher than that of proliferative cells during normal development of NSCs.

In order to preliminarily understand the mechanism of GA induced proliferation, we subsequently obtained that GA mainly exerted pharmacological activity through PDGF receptor and MAPK-ERK signal transduction pathway through RNA sequencing technology (RNA-Seq) analysis. The platelet derived growth factor (PDGF) is one of the receptor tyrosine kinase (RTK) family, located on the membrane surface of the PDGF family[23]. Its downstream mediates the regulation of multiple signaling pathways, such as PI3K-Akt pathway, MAPK kinase signaling cascade, and JAK-STAT pathway, etc. Therefore, we detecting the phosphorylation changes of key regulatory proteins in PI3K-Akt pathway and MAPK-ERK pathway by WB. Interestingly, we found that GA upregulated the phosphorylation level of ERK1/2 protein in the MAPK-ERK pathway rather than affecting the PI3K-Akt pathway. The specific proliferation mechanism of GA remains to be further studied.

Within MDHB and 8 polyphenol structural compounds, their structures were only one phenolic hydroxyl group or ester group different from MDHB, which is a natural compound small molecule with various neural activities. Hence, it is significant to compare the differences of these compounds and their neural activity. Through the analysis of these 9 compounds, we speculated that catechol structure was the main active group, which may be the key to the neurotrophic effect of the compound. When the three-dimensional configuration of the compound was changed, the pharmacological activity of the compound can vary markedly. Additionally, if the catechol structure is directly linked to the ester group, pharmacological activity may be generated to promote the differentiation of NSCs into neurons, while the introduction of 5-hydroxyl group may generate activity to promote the proliferation of NSCs. These groups can bind to different targets on the NSCs to produce diverse phenotypes. However, interestingly, although MG has both ester group and 5-hydroxyl group, the differentiated and proliferative activities are not desired. We suspect that the existence of multiple functional groups at the same time leads to the change of spatial binding pattern which generally resulting in a decrease of pharmacological activity. It is suggested that in the subsequent structural modification, we can try to change the binding mode by extend the carbon chain between ester group and benzene ring, so as to improve the efficacy. Furthermore, the drug activity was also closely related to the log*P* and hydrophobicity. When Lu and his team[24] studied the structure-activity relationship of GA and its derivatives, they found that the protective effect of GA on cell anti-oxidative stress injury not only depended on the scavenging ability of oxygen free radicals, but lay on the hydrophobicity of the structure. The higher the log*P* value, the more hydrophobicity the compound is, and the easier it is to penetrate the lipid bilayer into the cell to play its role. In our study, the structural formula and log P values of catechol and 9 tested compounds were analyzed **(Fig. 4A,B)**. Due to multiple phenolic hydroxyl molecules contained in GA, its stability and transmembrane ability are relatively weak, thus, it is highly likely that GA binds to the cell-membrane receptors rather than intracellular receptors to play a pharmacological role. Therefore, it deserves further study whether there is effective to maintain the stability of GA through preparing solution fresh when used or changing storage conditions, and to improve the bioavailability via making prodrugs, changing dosage forms or other useful ways. In future studies, we will further explore the genes inducing differentiation and proliferation to improve the mechanism study, and carry out structural transformation on the basis of the structure of GA to explore more possibilities of differentiation and proliferation activities on NSCs of catechol structural compounds.

**Fig. 4.**
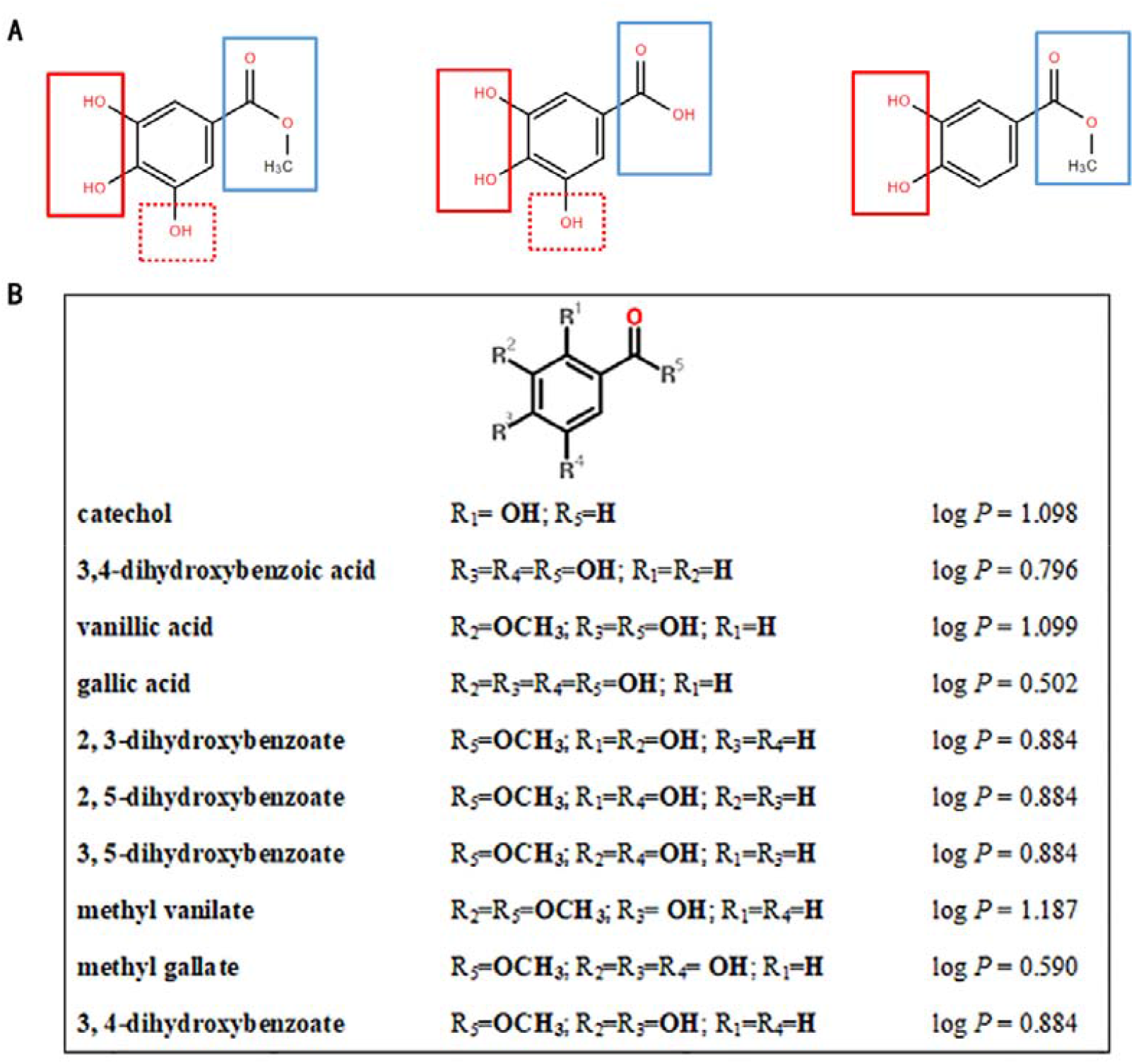
Structure-activity relationship of MDHB derivatives. **(A)** The structures of MG, GA, and MDHB. **(B)** The structures and log*P* values of catechol, MDHB, and its derivatives.

## Acknowledgements

We would like to thank the Medical Experimental Center of The Medical College of Jinan University.

## Abbreviations

AD: Alzheimer’s disease
Akt: protein kinase B
Aβ: amyloid beta protein
bFGF: basic fibroblast growth factor
CCK-8: cell counting kit-8
EGF: epidermal growth factor
ERK: extracellular regulated protein kinases
GA: gallic acid
GSK3β: glycogen syntheses kinase 3β
ICM: inner cell mass
iPS: induced pluripotent stem cells
MAPK: mitogen-activated protein kinase
MDHB: 3,4-dihydroxybenzoate
MG: methyl gallate
MV: methyl vanillate
NFT: neurofibrillary tangle
NMDA: N-methyl-D-aspartate
NSC: neural stem cell
PDGF: platelet derived growth factor
PDL: poly-D-lysine
PH3: phospho-histone H3
RPTKs: receptor protein tyrosine kinase
SGZ: subgranular zone
SVZ: subventricular zone

